# Soil solution in Swiss forest stands: a 20 year’s time series

**DOI:** 10.1101/2019.12.23.887042

**Authors:** Sabine Braun

## Abstract

The chemistry of the soil solution is influenced by atmospheric deposition of air pollutants, by exchange processes between the soil matrix and the soil solution and by processes between the rhizosphere and the soil. At sites of the Intercantonal Long-term Forest Observation Programme in Switzerland the soil solution has been monitored since 1998 in a number of forest plots growing from 9 to 47 sites in a wide range of soil conditions and air pollution impacts. The results show various site-specific developments of soil acidification. At sites with already advanced acidification (pH < 4.2), the acidification indicators remained rather stable at high levels, possibly due to the high buffering capacity of the aluminum buffer (pH 4.2 – 3.8). In contrast, in less acidified sites the acidification still progressed further which is reflected by e.g. the ongoing decrease of the base cation to aluminum ratio. Main driver of the acidification is the high N deposition which provokes cation loss and impedes sustainable nutrient balances for tree nutrition in the majority of plots examined. On an average for the years 2005-2017, N leaching rates were 9.4 kg N ha^-1^ yr^-1^, ranging from 0.04 to 53 kg N ha^-1^ yr^-1^. Three plots with high N input show very low N leaching, suggesting that N leaching may not always be a good eutrophication indicator. Both N deposition and N leaching have decreased since the year 2000 but the latter trend is partly also due to increased drought.

## 1 Introduction

Soil acidification is a serious threat for forest health ([1]). An increase in soil acidification, associated to atmospheric acid deposition, has been reported already in the 1980ies, for example in Germany ([2]) and Sweden ([3]). Due to mitigation measures, the deposition of acidifying substances in Europe, in particular of sulfur compounds, has decreased in recent years, but eutrophication due to the deposition of reactive nitrogen compounds remains an issue ([4], [5]). Between 1990 and 2017, the emissions of sulfur compounds in Switzerland have decreased by 85%, of oxidized nitrogen by 55% and of reduced nitrogen by 18% ([6])

Concerns on forest health in the 1980’s have led to the initiation of forest monitoring programs in which the monitoring of soil solution is crucial ([7]). The chemistry of soil solution is affected by atmospheric deposition, exchange processes between the solid and the dissolved phase in the soil, the nutrient uptake by the roots and rhizosphere processes ([8]). Stress indicators based on soil solution have been elaborated by expert groups under the International Cooperative Programme on Modelling and Mapping of Critical Loads and Levels and Air Pollution Effects, Risks and Trends (ICP Modelling und Mapping) of the Geneva Air Convention (CLRTAP) of the UNECE ([9], [10]). Important chemical criteria for evaluating the acidity of soil solution in forest ecosystems are the ratio between base cations and aluminum (BC/Al-ratio; [11]), the concentration of inorganic aluminum and the pH value. Critical thresholds have also been identified for base saturation (BS) of the solid phase and for the alkalinity and the acid neutralizing capacity of the soil solution.

Elevated N leaching from the rooting zone, mainly in the form of nitrate, induces acidification and the loss of base cations, impairing root development ([12], [13], [14]) and nutrient supply to plants ([2]). Sustainability calculations revealed that the loss of the cations Ca, Mg, and K due to nitrate leaching represents a greater risk for Swiss forest stands than cation exports with whole tree harvest ([15]). To protect forests from such negative effects, maximum tolerable values for N leaching have been set ([10]). Enhanced N leaching is not only important for the forest ecology but also for drinking water management as elevated nitrate concentrations pose a risk for human health.

N leaching is strongly linked to atmospheric N deposition ([16], [17]) and also C:N ratio in the forest floor has been shown to be a good predictor ([18]). A strong increase of N leaching has been shown at C:N ratios below 25. At higher ratios nitrogen is immobilized and nitrification is inhibited ([19]). High N leaching amounts have also been observed after disturbances such as tree cutting ([20]).

Aim of the present study is the description of the status and trends of the soil solution in the plots of the Intercantonal Forest Monitoring Program ([21]) and the evaluation of these findings in relation to forest health. Observed element concentrations and trends have been analyzed with explanatory variables and the results are presented in relation to critical limits and thresholds. The parameters measured in this monitoring program are based on the Guidelines on Reporting Monitoring and Modelling of Air Pollution Effects of the Geneva Air Convention ([22]).

## 2 Materials and Methods

### 2.1 Plots

The studied sites are part of the Intercantonal Forest Observation Program in Switzerland ([21]). In 1997 the first nine plots of the program (in total 189 plots) were equipped with suction cups, with samples being collected since 1998. More plots were added in subsequent years to give a current total of 47 plots with soil solution measurements (Table 1). A map of the plots with soil solution samplers is shown in Figure 1, site properties are listed in Table 1. The plots cover a wide range of forest soils. For description of the forest assessments and the soil and foliar analysis performed see [23].

**Table 1:** Site properties of the plots with soil water samplers. pH: pH(CaCl_2_) in the uppermost 40 cm of the soil, BS: base saturation in the uppermost 40 cm of the soil (%). CN: C:N ratio in the forest floor or the uppermost humus horizon. Prec: precipitation in mm, average 1981-2018. Leach: leaching in mm, calculated with the hydrological model Wasim-ETH ([24]), average 1981-2018. Tree species: Fa beech, Pic Norway spruce, Ab fir, La larch, Pin pine. Soil types: FAO classification. Weath: weathering rate in keq ha^-1^ a^-1^: calculations with SAFE ([25]) for the rooting zone (0-60 cm). Start: starting year of the soil solution measurements. Plot with denomination “storm”: plot cleared in 1999 during the gale “Lothar”.

| Site | abbr. | altitude | prec | leaching | species | pH | BS | CN | soil type | weight | start |
| --- | --- | --- | --- | --- | --- | --- | --- | --- | --- | --- | --- |
| Aarwangen | AW | 470 | 1140 | 482 | Fa | 3.99 | 10 | 14.5 | Dystric Cambisol | 1.31 | 2002 |
| Aeschau | AU | 940 | 1512 | 783 | Ab Pic (Fa) | 3.67 | 20 | 26.0 | Dystric Arenosol | 0.45 | 1997 |
| Aeschi | AI | 510 | 1160 | 472 | Fa Pic | 3.87 | 15 | 21.2 | Haplic Luvisol | 1.24 | 1998 |
| Allschwil | AL | 350 | 896 | 153 | Pic | 4.31 | 88 | 14.0 | Haplic Luvisol |  | 2006 |
| Bachtel | BAB | 1030 | 1825 | 1093 | Fa | 3.93 | 36 | 15.6 | Chromic Luvisol | 4.62 | 1999 |
| Bachtel | BA | 1040 | 1770 | 998 | Pic | 4.01 | 7 | 24.8 | Chromic Luvisol | 0.90 | 1997 |
| Beromünster | BE | 640 | 1220 | 321 | Pic | 5.00 | 90 | 23.1 | Gleyic Cambisol | 7.06 | 2016 |
| Bonfol | BO | 450 | 1091 | 417 | Fa | 4.26 | 18 | 20.3 | Dystric Cambisol | 0.67 | 2004 |
| Braunau | BRAU | 710 | 1253 | 400 | Pic | 4.05 | 55 | 19.8 | Haplic Luvisol |  | 2006 |
| Breitenbach | BB | 460 | 1111 | 346 | Fa | 4.53 | 91 | 14.3 | Haplic Luvisol | 0.85 | 2003 |
| Brislach beech | BRB | 435 | 1041 | 378 | Fa | 4.09 | 25 | 13.3 | Haplic Luvisol | 0.88 | 2000 |
| Brislach spruce | BR | 435 | 1042 | 258 | Pic | 3.93 | 12 | 23.3 | Haplic Luvisol | 0.84 | 1997 |
| Bürglen | BUR | 640 | 1582 | 572 | Pic | 4.77 | 99 | 22.2 | Cambisol | 0.36 | 2016 |
| Buswil | BU | 600 | 1195 | 388 | Pic | 3.78 | 3 | 18.9 | Haplic Luvisol | 0.99 | 2004 |
| Diessenhofen | DI | 520 | 942 | 290 | Pic | 3.7<br>7 | 16 | 20.<br>8 | Dystric Cambisol |  | 200<br>6 |
| Frienisberg | FR | 725 | 120<br>9 | 542 | Fa Pic | 3.9<br>0 | 21 | 21.<br>2 | Dystric Arenosol | 0.64 | 199<br>7 |
| Gelfingen | GE | 540 | 113<br>5 | 451 | Fa | 6.5<br>5 | 10<br>0 | 21.<br>9 | Calcaric Cambisol | 1.59 | 201<br>6 |
| Giswil | GI | 540 | 130<br>6 | 479 | Fa | 5.8<br>6 | 10<br>0 | 19.<br>5 | Calcaric Cambisol | 10.8<br>4 | 201<br>6 |
| Grenchenberg | GB | 1220 | 151<br>1 | 961 | Fa Pic | 5.6<br>4 | 10<br>0 | 15.<br>1 | Calcaric Cambisol | 19.0<br>5 | 199<br>7 |
| Grosswangen | GW | 600 | 111<br>4 | 320 | Pic | 3.5<br>2 | 14 | 21.<br>9 | Stagnic Acrisol | 1.25 | 201<br>6 |
| Habsburg K | HA | 430 | 107<br>2 | 308 | Fa | 4.1<br>7 | 16 | 17.<br>1 | Haplic Luvisol | 0.84 | 200<br>4 |
| Hinwil | HI | 650 | 145<br>6 | 619 | Pic | 5.1<br>2 | 95 | 15.<br>4 | Eutric Cambisol | 1.33 | 200<br>2 |
| Le Châtelard | LC | 1010 | 165<br>4 | 811 | Pic | 3.7<br>4 | 20 | 29.<br>3 | Gleyic Cambisol | 1.53 | 200<br>6 |
| Lurengo spruce | LUB | 1620 | 178<br>6 | 1098 | Pic Pin La | 3.9<br>0 | 28 | 26.<br>2 | Dystric Arenosol |  | 199<br>9 |
| Möhlin | MO | 290 | 103<br>4 | 267 | Pic | 3.7<br>9 | 12 | 17.<br>5 | Haplic Luvisol | 1.22 | 199<br>8 |
| Muri beech | MUB | 490 | 111<br>0 | 340 | Fa | 4.0<br>0 | 24 | 18.<br>3 | Haplic Luvisol | 0.56 | 199<br>9 |
| Muri spruce | MUF | 490 | 110<br>4 | 278 | Pic | 3.8<br>8 | 10 | 26.<br>5 | Dystric Cambisol | 0.74 | 200<br>1 |
| Muri storm | MU | 490 | 110<br>4 | 588 | Pic | 4.0<br>8 | 23 | 18.<br>9 | Haplic Luvisol | 0.62 | 199<br>7 |
| Muttenz | MUU | 375 | 912 | 228 | Fa | 4.06 | 41 | 15.7 | Stagnic Luvisol | 0.50 | 2004 |
| Oberschrot | OS | 950 | 1340 | 541 | Fa Pic | 3.61 | 11 | 17.2 | Gleyic Stagnic Cambisol |  | 2006 |
| Olsberg | OL | 380 | 998 | 240 | Fa | 4.06 | 20 | 15.4 | Dystric Planosol | 0.48 | 2004 |
| Pratteln | PR | 415 | 966 | 339 | Fa | 5.15 | 100 | 12.4 | Chromic Luvisol | 2.24 | 2002 |
| Rafz | RAF | 540 | 995 | 315 | Pic | 4.18 | 16 | 19.0 | Haplic Luvisol | 0.70 | 2004 |
| Riehen | RI | 470 | 1005 | 402 | Fa | 6.41 | 100 | 13.3 | Haplic Luvisol | 1.26 | 2002 |
| Rünenberg | RU | 590 | 1017 | 245 | Fa | 4.13 | 35 | 17.2 | Haplic Luvisol | 0.68 | 2002 |
| Sagno | SA | 770 | 1782 | 943 | Pic | 3.83 | 25 | 21.8 | Eutric Cambisol | 0.41 | 1999 |
| Scheidwald | SW | 1170 | 1500 | 547 | Pic | 3.41 | 7 | 27.9 | Dystric Gleysol | 0.66 | 2008 |
| Sempach | SE | 550 | 1139 | 450 | Fa | 3.71 | 39 | 21.6 | Gleyic Luvisol | 2.32 | 2016 |
| Stans | ST | 560 | 1437 | 924 | Fa | 6.40 | 100 | 17.4 | Calcaric Cambisol | 28.30 | 2016 |
| Wangen | WG | 500 | 1143 | 450 | Fa Pic | 3.88 | 24 | 23.3 | Chromic Luvisol |  | 2008 |
| Wangen SZ | WSZ | 470 | 1536 | 634 | Fa | 4.43 | 95 | 14.8 | Luvisol | 1.77 | 2016 |
| Wengernalp | WA | 1870 | 1605 | 922 | Pic | 3.53 | 28 | 14.2 | Podzol | 0.15 | 1997 |
| Winterthur | WI | 530 | 1178 | 465 | Pic | 5.25 | 97 | 16.0 | Vertisol | 18.46 | 2003 |

Table 1: Site properties of the plots with soil water samplers.
| Site | abbr. | altitude | precip | leaching | species | pH | BS | CN | soil type | weathering | start |
| --- | --- | --- | --- | --- | --- | --- | --- | --- | --- | --- | --- |
| Zofingen | ZO | 540 | 1130 | 370 | Fa Pic | 4.00 | 17 | 17.9 | Haplic Luvisol | 0.92 | 2004 |
| Zugerberg HG | ZBB | 980 | 1569 | 900 | Fa Pic | 4.20 | 37 | 19.8 | Eutric Cambisol | 0.63 | 1999 |
| Zugerberg (N-exp.) | ZB | 940 | 1457 | 821 | Fa | 3.91 | 100 | 18.5 | Dystric Cambisol | 0.45 | 1997 |
| Zugerberg VG | ZV | 900 | 1457 | 550 | Pic | 3.62 | 24 | 20.2 | Dystric Cambisol | 1 | 2002 |

**Figure 1:**
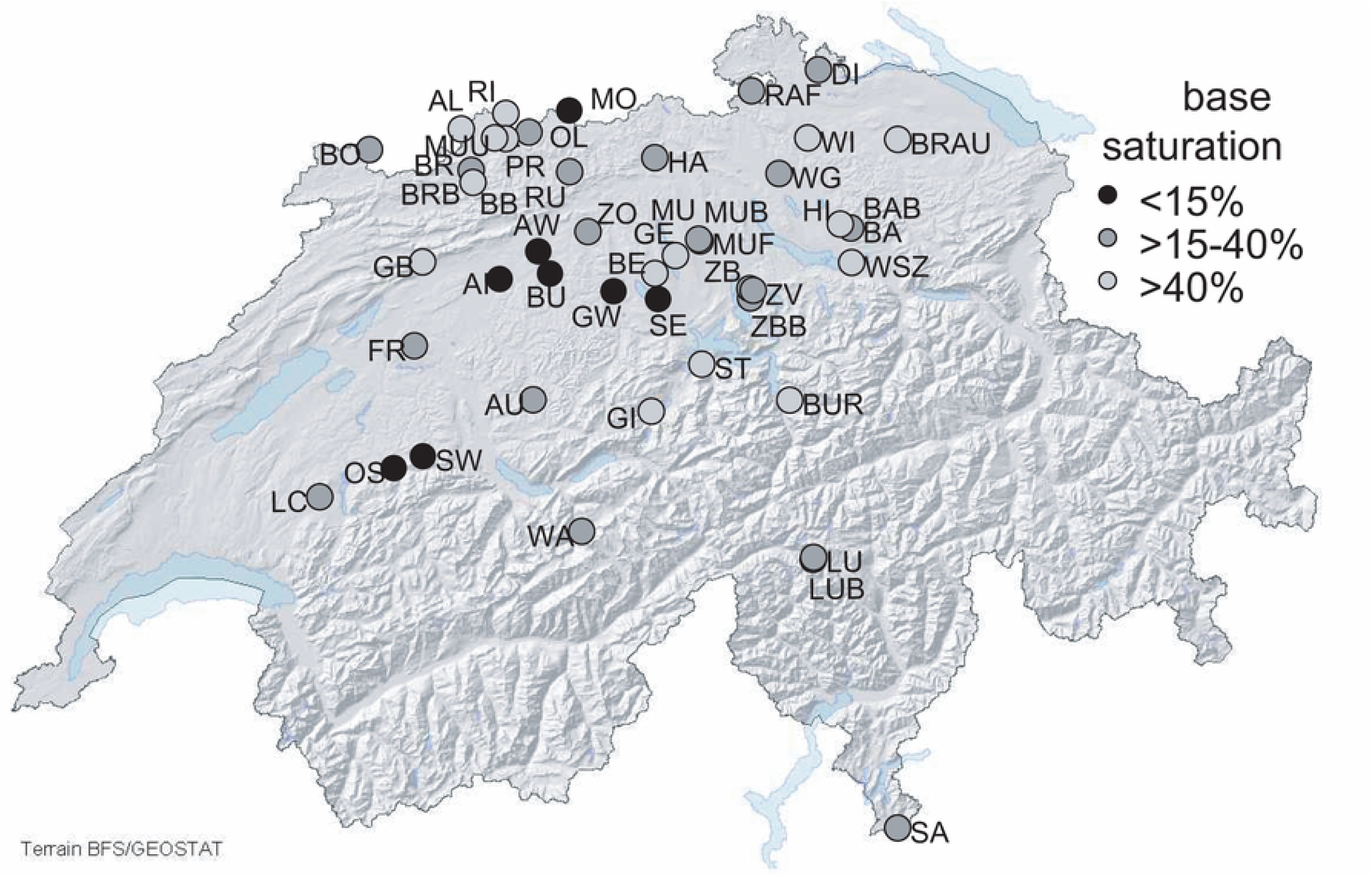
Forest plots with soil solution samplers, grouped according to base saturation of the topsoil (average 0-40 cm). Plots sampled in 2018.

Tree mortality rates and the rates of removed trees were derived from annual observations. Changes were related to the population of the previous year and lagged for up to four years. Mortality and tree removal were combined to one variable further called “tree removal”.

### 2.2 Soil solution

Per site and soil depth 8 replicate samplers were installed in the upper soil and 5 in the subsoil. Detailed site specific information can be obtained from a project report ([26]). Soil solution was sampled once a month. After collection, samples from the same site and depth were pooled. pH, conductivity and alkalinity were measured immediately after sampling. Samples were then kept frozen until further analysis. For cation analysis the samples were acidified with 0.5 ml HNO_3_ conc in 10 ml solution prior to freezing, for anion analysis they were filtered through a 0.45 µm membrane filter.

The following parameters were measured in the soil solution:

- pH: Metrohm pH-meters 716 and 809, with Metrohm Aquatrode (part No. 6.0257.000)
- conductivity: Metrohm conductivity meters 712 and 856, with Metrohm cell 6.0916.040.
- Ca, Mg, Al, Mn; Atomic absorption photometry (Varian 640)
- K, Na: flame photometry (Varian 640)
- inorganic Al: Al was measured before and after passing the samples through an ion exchanger (0.5 ml IC-H, Alltech 30264).
- NH_4_^+^: photometric determination with indophenol blue ([27])
- NO_3_^-^, SO_4_^2-^, Cl^-^: ion chromatography with suppressed conductivity (Dionex GP50 pump, ED50 electrochemical detector and AS3500 autosampler)
- alkalinity: titration with HCl to a pH of 4.35 (Metrohm 809).
- UV absorption at 280 nm for calculation of dissolved organic carbon (DOC) according to [28](2001)

Quality control was achieved by calculation of the ion balance, by comparison of measured and calculated conductivity ([29,30]) and by analysis of reference samples distributed once a year by the Norwegian Institute for Air Research (NILU).

The determination of organic aluminum was started in 2005. In order to get a homogenous time series also for older data, the average proportion of organic aluminum to total aluminum was calculated for each soil layer. This proportion was then applied to data from 1998 to 2005. It varied between 50% for the uppermost soil water samplers and 25% in the lowest ones (Figure_Supplementary 2).

The amount of leaching water per hectare was calculated using the hydrological model Wasim-ETH ([24]) using the sites soil characteristics (pF curve, texture) and daily meteorological data interpolated for each site ([23]). Leaching fluxes calculated for each sampling period were multiplied with concentrations to calculate element fluxes.

#### 2.2.1 Critical limits in soil solution

The molar ratio between base cations (BC = Ca^2+^, Mg^2+^, K^+^) to aluminum (Al^3+^), the BC/Al ratio, in the soil solution is an important criterion for the status of soil acidification. It has been shown to be closely correlated with growth and vitality parameters of the vegetation ([11]). Initially, a limit of 1 has been set for the BC/Al ratio ([9], [11]). As this limit is not necessarily sufficient to protect forests, Ouimet et al. (2006) suggested a BC/Al limit of 10 for calculations of critical loads in Canada. Therefore the critical limits of the BC/Al ratio were revised according to better knowledge ([31]) In Switzerland a limit of 7 has recently been applied ([32]).

Other critical limits for the soil solution are a pH of 4 and a Al concentration of 0.2 eq m^-3^ ([10]). The Acid Neutralizing Capacity (ANC), an indicator of susceptibility against acidification, is defined as sum of the base cations Ca, Mg, K and Na minus the sum of the anions nitrate, sulfate and chloride. It relates the two criteria Al and proton concentration, given a pK_Gibbsit_ of 8.04([33]):

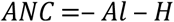

When the above equation is solved for Al_crit_ of 0.2 eq m^-3^ and pH 4, a maximum allowable leaching of alkalinity from the rooting zone of −300 µeq l ^-1^ is resulting ([34]).

Monthly soil solution data were compared with the critical limits listed above. In order to avoid sensitivity to single outliers, an exceedance was indicated when more than 1% of the values exceeded the limit.

Eutrophication effects of N input can be evaluated by concentrations and total amounts of N-leaching. Critical limits have been set accordingly. A concentration of >0.2 mg N l^-1^ in soil solution has been related to changes in ground vegetation and in tree nutrition ([10]). For temperate deciduous forests, a threshold for a total annual N leaching amount of 2-4 kg N ha^-1^ yr^-1^ has been set to avoid excessive base cation leaching and acidification. Absolute limits for N leaching are especially important for areas with a high precipitation since limits based on concentrations in soil solution will lead to high losses of cation nutrients under these conditions, leading to a loss of base saturation in the soils.

While the above mentioned acidity indicators refer to soil solution, an indicator widely used in forestry is the base saturation (BS) of the soil solid phase ([35]).

### 2.3 Weathering rates

Weathering rates were calculated using the model SAFE ([25]) for a subset of monitoring plots. The calculations based on measured mineralogy of soil samples ([36]). The results were summed up per layer down to a depth of 60 cm except when dense rooting was observed also in deeper layers. This differentiation has been based on correlation analysis between soil chemistry and foliar analysis suggesting that below 60 cm not much nutrient uptake occurs ([15]).

### 2.4 Atmospheric Deposition

Data on atmospheric deposition of reactive nitrogen compounds and base cations were received from the Swiss Federal Office for the Environment ([37]). N deposition was modelled in a spatial resolution of 0.1 ha; deposition of base cations according to [38] with a spatial resolution of 2 km. Comparison of the modelled N deposition with measurements showed good agreement ([39]) except for plots in Southern Switzerland where the import of N compounds via the air from Italy is more difficult to account for.

### 2.5 Statistics

In order to analyze the development of the BC/Al ratio against various predictors, moving time windows of 5 years were formed. The development within these time windows was analyzed using a mixed regression with plot as random variable (R, package lme4, [40]). The following predictors were used:

- BC/Al at the start of the 5 year period
- Soil solution pH at the start of the 5 year period
- Base saturation of the soil in the corresponding soil horizon (sampling of soil solid phase in 2005)
- Weathering rate
- Proportion of coniferous trees in the plot
- Modelled N deposition
- Organic carbon in the corresponding soil horizon
- C:N and N:P ratio in the forest floor
- Clay content
- Depth (binary variable coded as ≤70 and >70 cm).

Explanatory variables for N leaching were analyzed based on annual averages. A mixed regression with plot and year as random variable was used. The following predictors were included in the regression model:

- Modelled N deposition
- C:N in the uppermost soil horizon
- Water holding capacity cumulated over the uppermost 100 cm
- Potential evapotranspiration (annual sum).
- Annual minimum of site water balance.
- Rate of seepage water.
- Tree removal: current year and lagged by up to 4 years
- Shrub cover of the plot (vegetation survey)
- Proportion of coniferous trees in the plot
- Altitude

Predictors were selected backwards using the Akaike Information Criterion (AIC) as indicator which should be minimized. When the number of predictors had to be reduced to avoid oversaturation of the model, also the Bayes Information Criterion (BIC) criterion was used as indicator. Residuals were checked for normal distribution, homoskedasticity and outliers by diagnostic plots. Regression plots were produced using the R functions ggpredict (library ggeffects, [41]) and ggplot (library ggplot2, [42]). The former extracts predictions from a multivariate model with all other predictors set at their mean and a 95% interval.

## 3 Results

### 3.1 Acidification

#### 3.1.1 Acidification status, comparison with thresholds

Table 2 lists the exceedance of various acidity limits set under the Geneva Air Convention ([10]) or suggested by [43]. The BC/Al values are lower than 1 in 27% of the plots (in at least one layer) and lower than 7 in 71% of the plots.

**Table 2:**
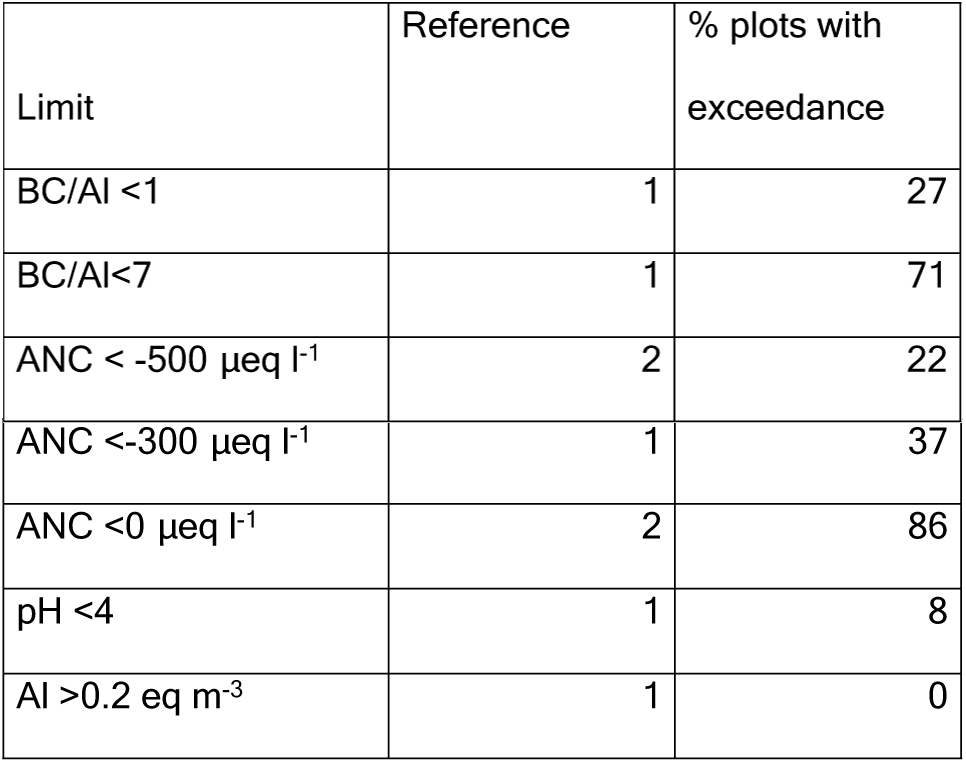
Frequency of exceedance of various acidity limits in the years 2013-2018. Reference to limits: 1) [10], 2) [43]).

The BC/Al ratio is closely related to the base saturation in the corresponding soil layer and somewhat less with the pH(CaCl_2_) of the solid phase (Figure 2). These relationships allow – within certain limits – to link BC/Al ratios with base saturation values which are more often used as acidity indicator in forestry. For BS=20 the regression function predicts a BC/Al ratio of 12; for BS=40 a BC/Al of 51. The less clear relationship of the BC/Al ratio with pH(CaCl_2_) can be attributed to the high buffer capacity of the aluminum buffer (pH 3.8 – pH 4.2; 150 kmol H^+^ per % clay), reflected by the cluster of points around pH 4.

**Figure 2:**
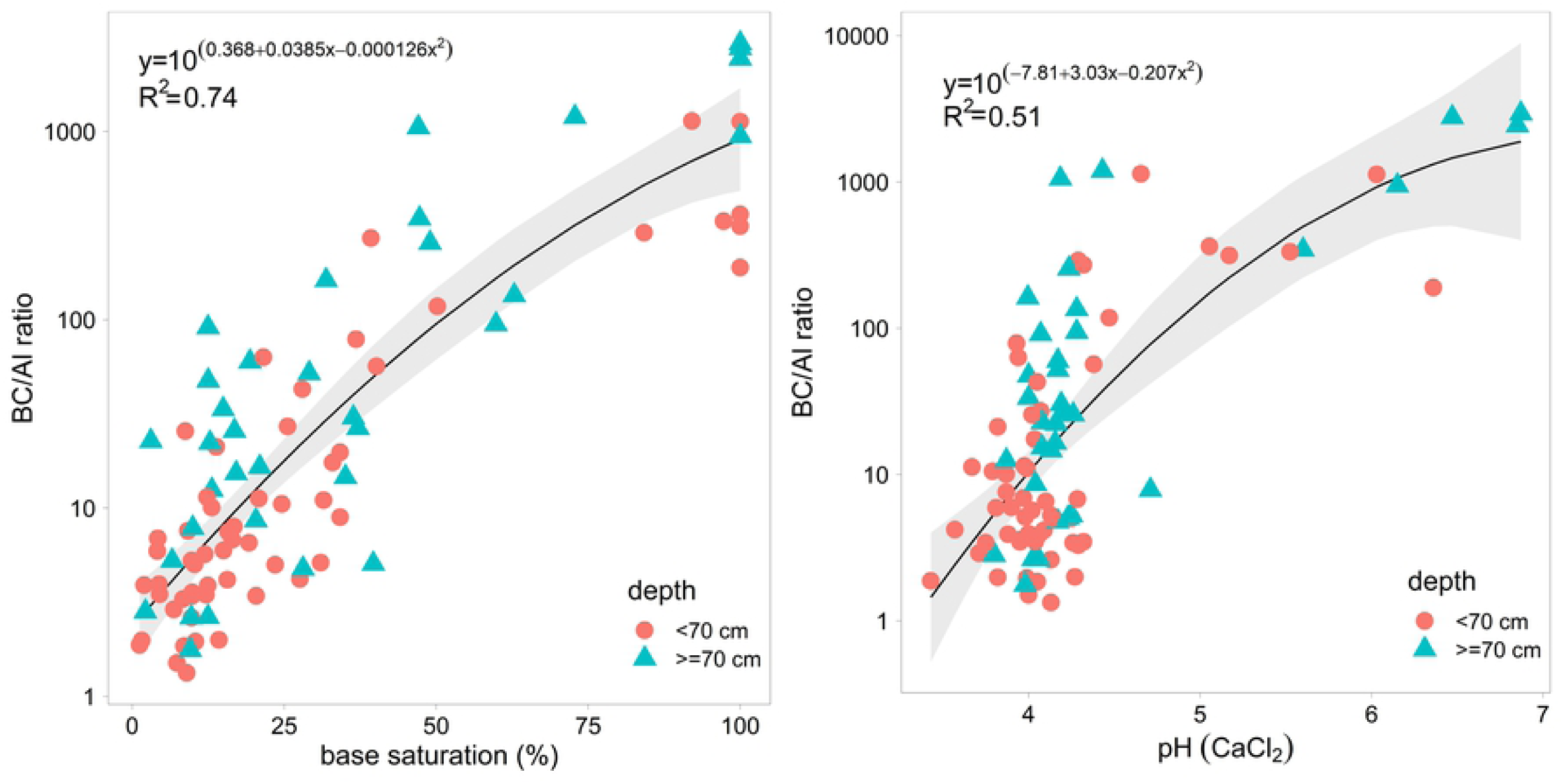
Relation between the BC/Al ratio of the soil solution with BS and pH(CaCl_2_) of the corresponding horizon.

#### 3.1.2 Leaching of base cations

The leaching of base cations is relevant for the assessment of acidification and the evaluation of the nutritional sustainability of a forest site. The relation between the input from weathering and deposition on the one hand and leaching on the other hand was clearly significant only for Ca. Figure 3 shows this relation between leaching and either weathering or total input (weathering + deposition). In 35 out of 41 plots (85%), Ca leaching exceeded the Ca input by weathering. When atmospheric deposition of base cations is added to weathering, 22 plots (=54%) still have a negative balance. The highest Ca losses were observed in weakly buffered plots in the silicate and exchange buffer range (WI, HI, PR, BB) as well as in a plot with very high N input (SA). Leaching was most correlated with weathering rate and slightly with the water holding capacity of the soil (total variance explained 58%) while neither N deposition nor species composition of the tree layer were significant predictors.

**Figure 3:**
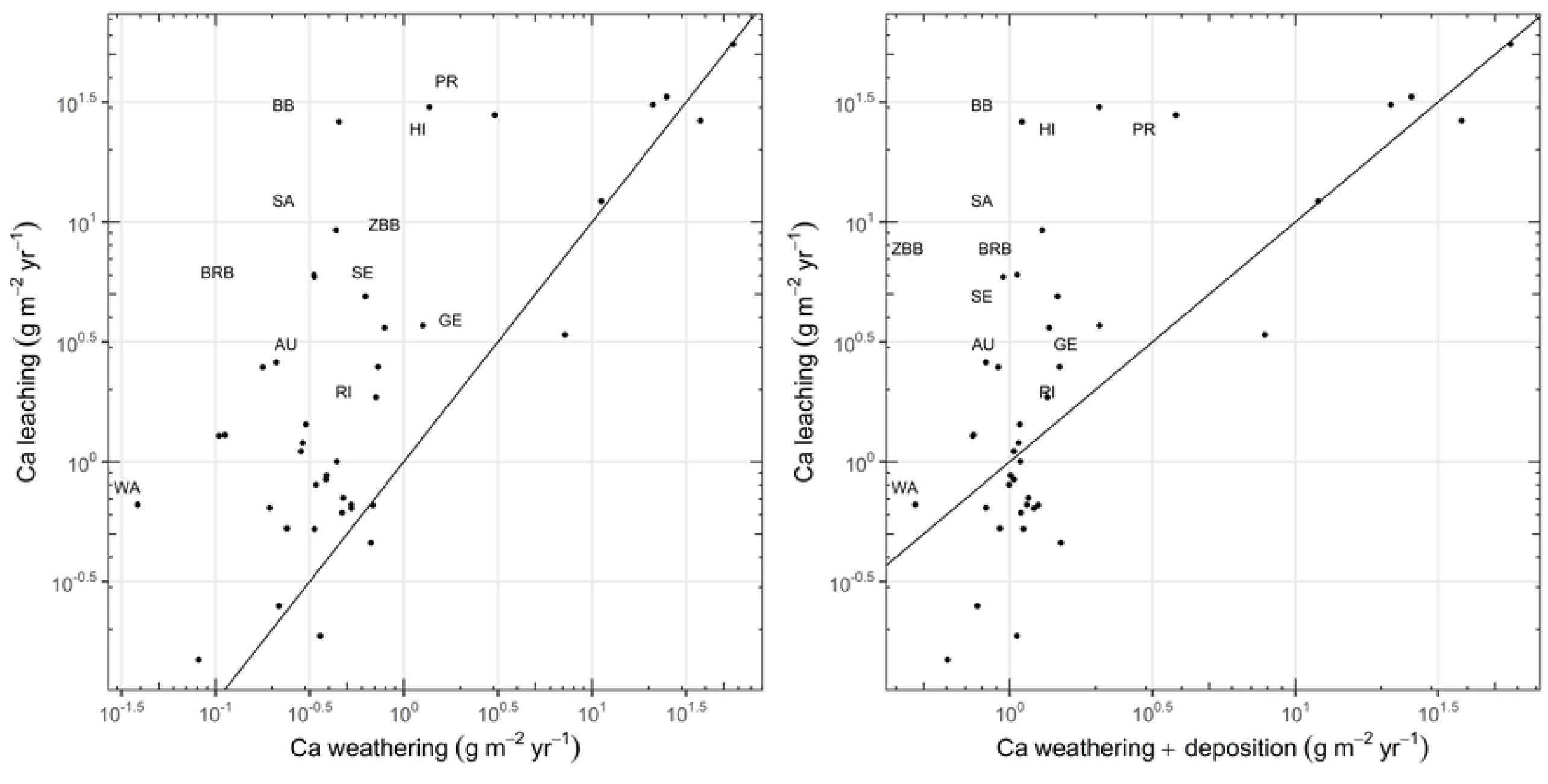
Relation between the leaching of Ca^2+^ at a depth of 60-80 cm (average 2005-2018) and the weathering rate of base cations cumulated over the uppermost 60 cm. Left: relation with weathering rate only, right: with the sum of Ca weathering and Ca deposition (model [36], for Southern Switzerland [44]). Site abbreviations see Table 1. The line is the 1:1 line.

#### 3.1.3 Development of acidification

On an average of all plots, the BC/Al ratio decreased significantly over time. Figure 4 shows this development for soils grouped according to the base saturation in the uppermost 40 cm. Even in base rich soils there is a clear decrease of the BC/Al ratio, i.e. a clear increase in acidification. The decrease is strongest in the uppermost soil. Below the rooting zone the changes are weaker but still clear.

**Figure 4:**
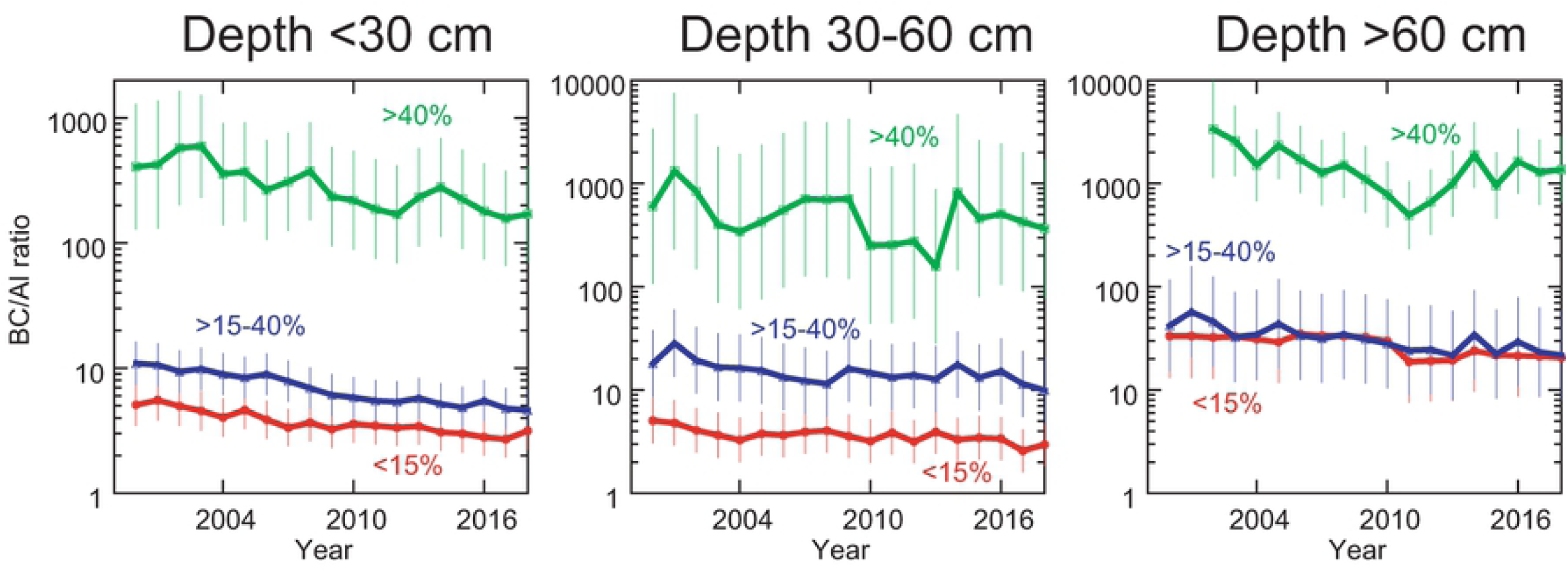
Development of the BC/Al ratio in soils of different BS. Graph corrected for varying plot number in different years by using estimates from a mixed regression with plot as factor. Error bars: 95% confidence interval, extracted from the mixed regression with ggpredict (R package ggeffects, [41]).

The decreasing trends are getting weaker the stronger the soils are acidified. To examine this association, the changes of BC/Al ratio within five years were analyzed in relation to the initial status of the sample in a moving window analysis. The resulting regression (Table 3) supports the hypothesis that the trends are related to acidification status, and Figure 5 illustrates the observed association with predicted values. Significant predictors for the trend were the initial BC/Al ratio and the initial pH, the BS at the corresponding soil depth and depth as a binary variable (<70, >= 70). Predicted changes of log BC/Al in Figure 5 below zero correspond to an expected decrease of the BC/Al ratio during the following five years. This means, that a decrease of BC/Al is expected when the BS is below 48%, the pH below 5.9 or the BC/Al ratio above 16.3. The value of pH 5.9 corresponds well with the lower limit of the Ca buffer range (pH 6.2), according to the equilibrium of CaCO_3_ with H_2_CO_3_ in soil ([45], [46]). These values may be regarded as “threshold” for acidification under the current deposition situation. No interpretation was found for the residuals of this regression, i.e. the unexplained part of the trend.

**Table 3:**
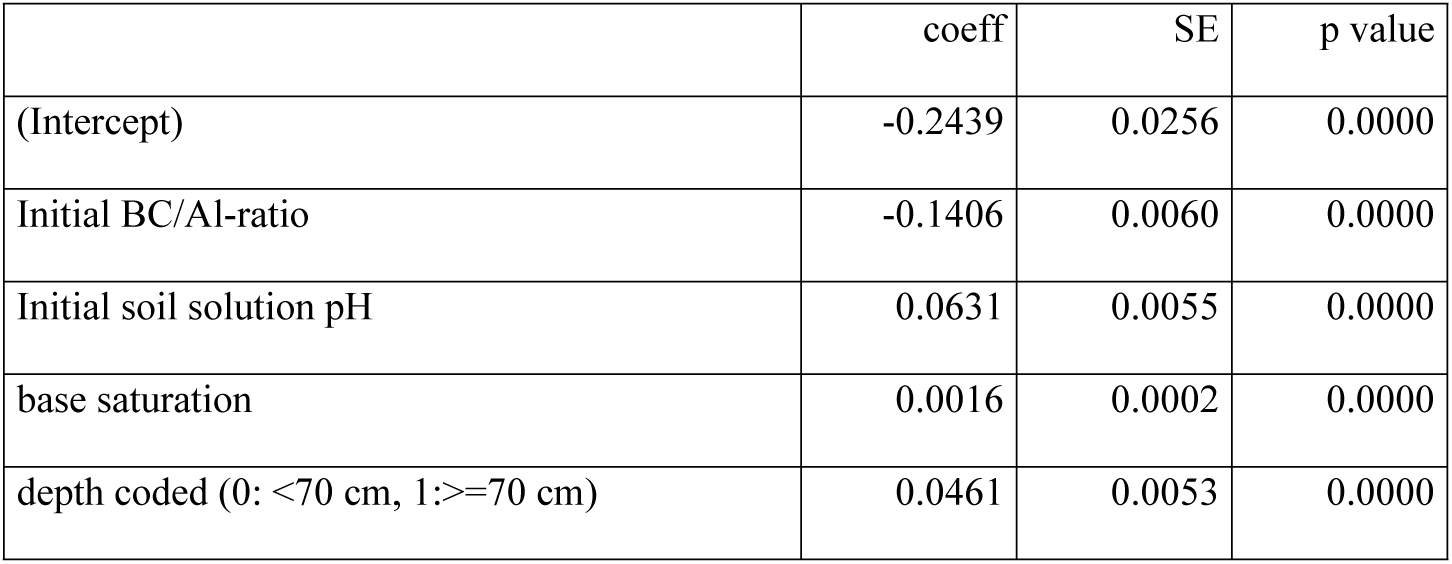
Speed of the change of BC/Al in relation to various parameters. Dependent variable: Difference in log transformed BC/Al during five years. The regression explains about 36% of the variance of the trend.

|  | coeff | SE | p value |
| --- | --- | --- | --- |
| (Intercept) | -0.2439 | 0.0256 | 0.0000 |
| Initial BC/Al-ratio | -0.1406 | 0.0060 | 0.0000 |
| Initial soil solution pH | 0.0631 | 0.0055 | 0.0000 |
| base saturation | 0.0016 | 0.0002 | 0.0000 |
| depth coded (0: $<70$ cm, 1: $\geq 70$ cm) | 0.0461 | 0.0053 | 0.0000 |

**Figure 5:**
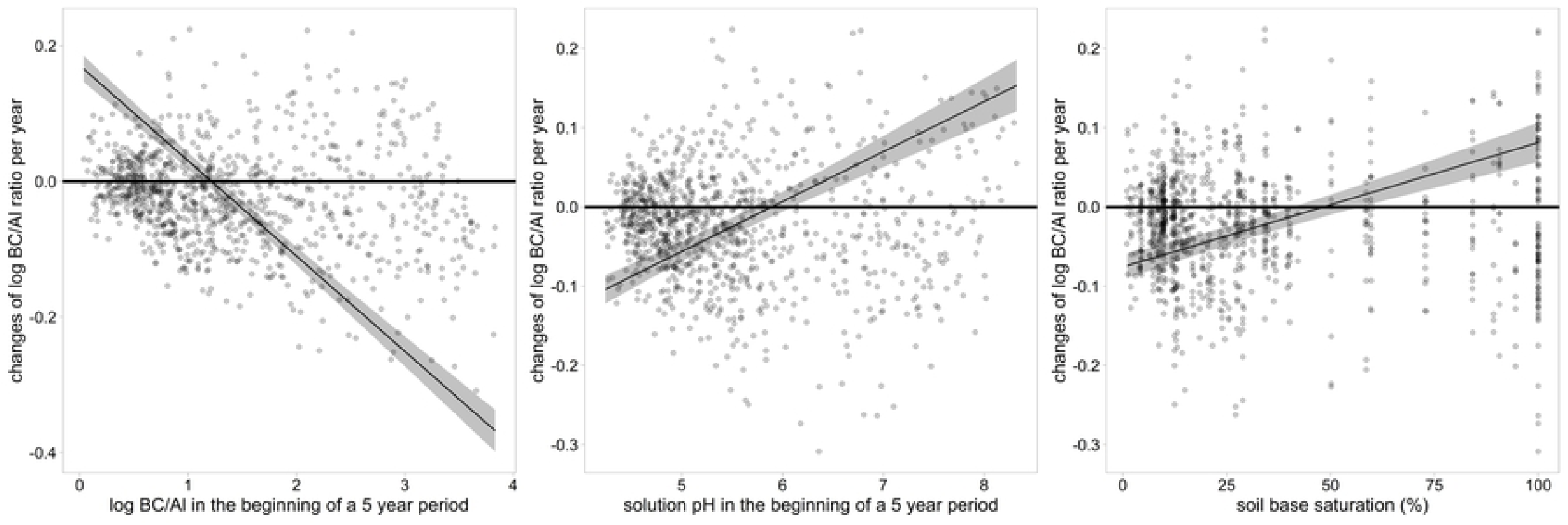
Speed of the change in BC/Al in relation to the predictors in Table 3. Values predicted from the regression with all other predictors to their mean. Negative changes signify an expected decrease in BC/Al. The linearity of the relations was tested using polynomial functions.

#### 3.1.4 Nitrogen leaching

Nitrogen leaching decreased between 1998 and 2018 (Figure 6). The annual nitrogen leaching rates in the years 2005-2018 were on an average 9.5 kg N ha^-1^ yr^-1^. The proportion of plots exceeding the leaching limits set under the Geneva Air Convention [10] decreased from 83 % (1998, 12 plots) to 34% (2018, 47 plots). For reasons of better comparability, Table_Supplementary 1 lists the average N leaching over the time period 2005-2018. The two plots with the highest average leaching rates have very different properties. In southern Switzerland (plot SA), a leaching rate of 52 kg N ha^-1^ yr^-1^ results from a high N deposition, leading to an average nitrogen concentration of 6.7 mg N l^-1^, and a high output with the seepage water (>900 mm yr^-1^). The second plot, in Northwestern Switzerland (plot AL), is characterized by extremely high N concentrations in soil water (average 30 mg N l^-1^) and a low leaching water flux, resulting in an average leaching rate of 55 kg N ha^-1^ yr^-1^. Despite high nitrogen deposition and low base saturation three plots have almost negligible N leaching (GW, SW, BU). Interestingly enough these plots stand out with a very high crown transparency (proportion of Norway spruce with >25% transparency in 2016, 2017 and 2018 was: 81%, 74% and 43%, respectively, while the average proportion in all 76 Norway spruce plots was 23.4%).

**Figure 6:**
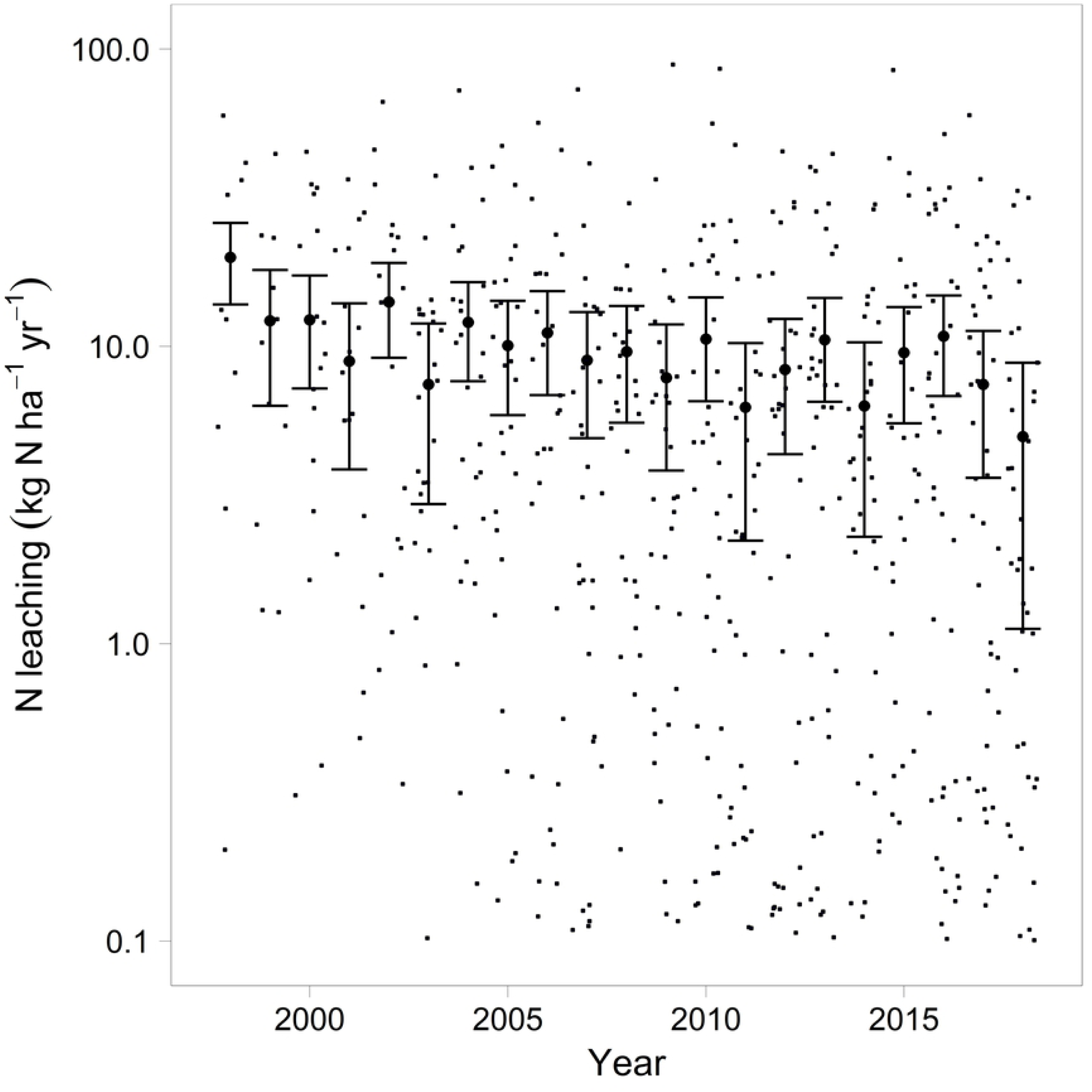
Development of N leaching since 1998. The decrease with time is significant with p<0.001. Thick dots and error bars (95% confidence intervals) are estimates corrected for the varying number of plots per year (predictions from mixed regression). Small dots are raw data.

The relation between N leaching rates and various site predictors was examined in a data set with annual climate data, modelled N deposition including a time trend and with site characteristics. N deposition, drought, tree removal and water holding capacity were significant predictors for N leaching (Table 4, Figure 7). Drought predictors included potential evapotranspiration, site water balance and the amount of leaching water. The effect of tree removal was largest in the year after removal (lag 1, Table 4). In the year of the removal (lag 0) and four years after removal (lag 4) it was not significant but the inclusion of the year of removal slightly improved the model. For graphical illustration the annual removal rates were combined to a weighted average based on the model coefficients in Table 4 for lag 0, 1, 2 and 3 (Figure 8). N leaching was reduced in dry years (Figure_Supplementary 5) and on soils with a high water holding capacity (Figure_Supplementary 6). C:N ratio in forest floor was not a significant predictor for N leaching.

**Table 4:** Regression analysis of N leaching with annual data. Dependent variable: N leaching in kg N ha^-1^ yr^-1^, log transformed.

|  | coeff | SE | p-value |
| --- | --- | --- | --- |
| (Intercept) | 4.185 | 1.683 | 0.0129 |
| N deposition | 0.101 | 0.021 | 0.0000 |
| removed trees current year (lag 0) | 0.954 | 0.581 | 0.1008 |
| removed trees previous year (lag 1) | 2.588 | 0.578 | 0.0000 |
| removed trees two years before (lag 2) | 1.956 | 0.583 | 0.0008 |
| removed trees three years before (lag 3) | 1.085 | 0.494 | 0.0280 |
| potential evapotranspiration | -0.003 | 0.001 | 0.0006 |
| minimum site water balance | -2.092 | 0.804 | 0.0092 |
| leaching water*1000 | 1.066 | 0.259 | 0.0000 |
| water holding capacity 0- 100 cm (mm) | -0.014 | 0.004 | 0.0011 |

**Figure 7:**
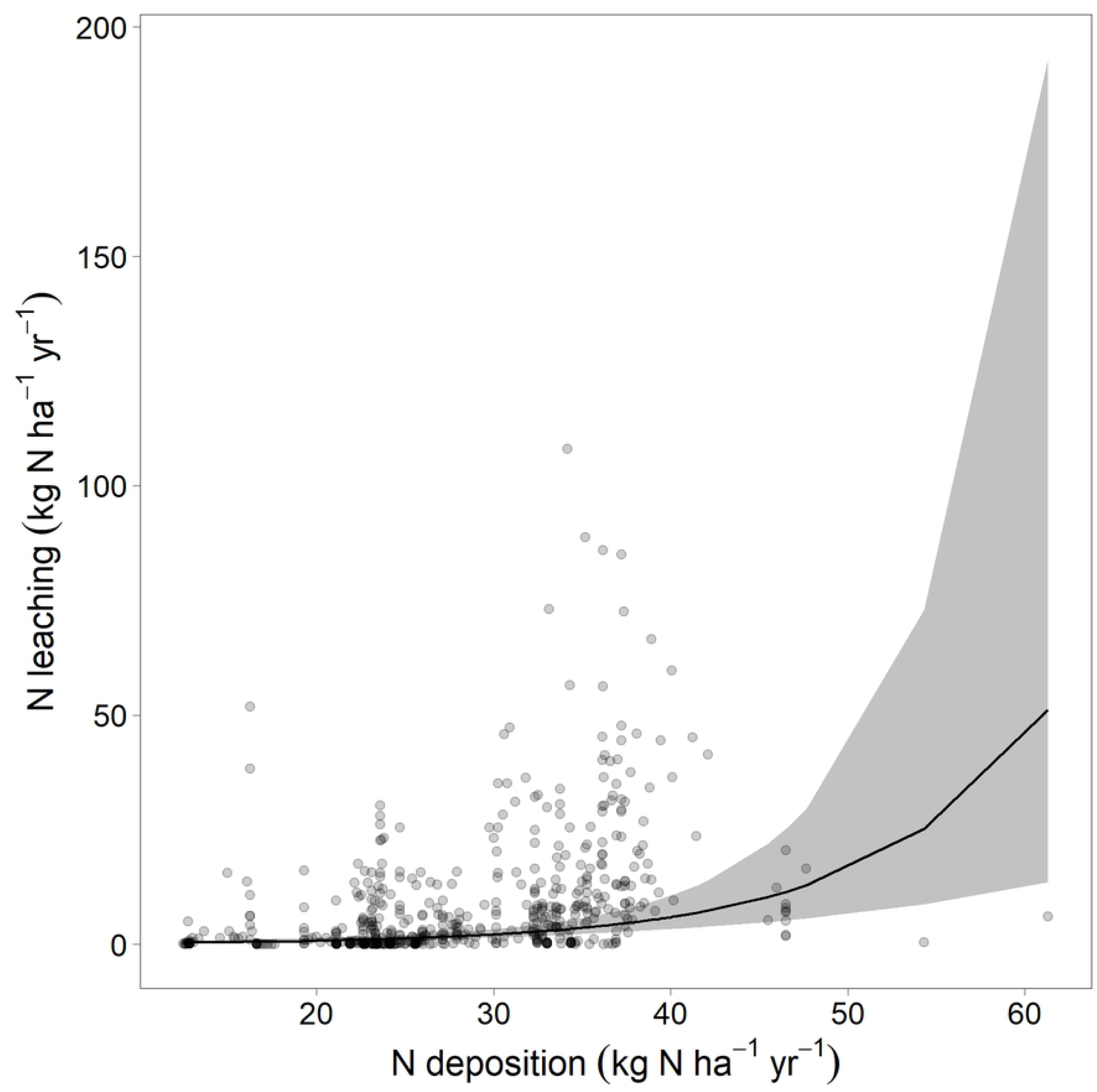
Illustration of the relation between N leaching and N deposition as calculated with the regression model presented in Table 4. Points are annual leaching rates per plot. The two points with high N leaching at 17 kg N ha^-1^ yr^-1^ input are the result of a bark beetle attack.

**Figure 8:**
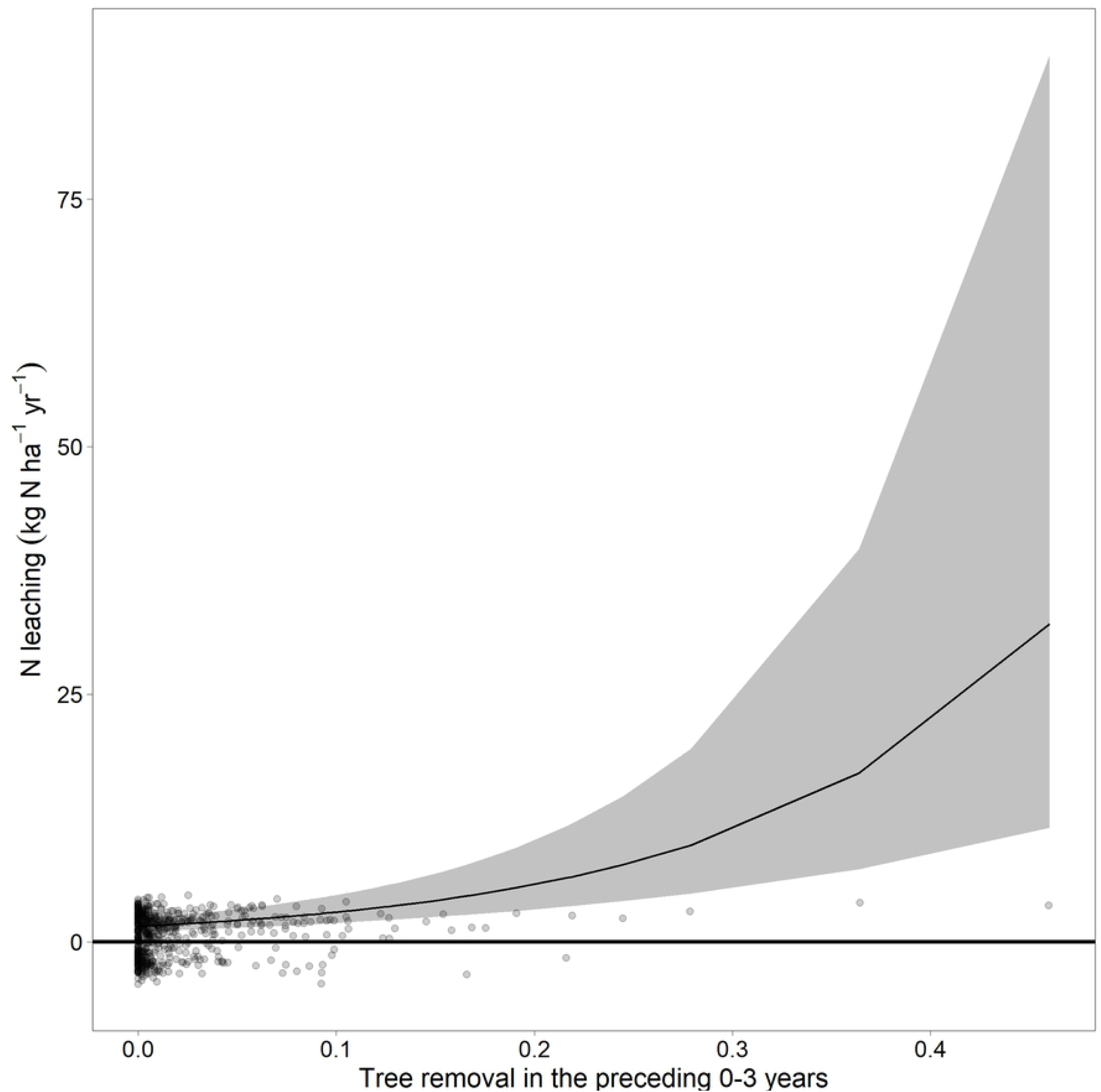
Relation between N leaching and tree removal, averaged over 0-3 years according to the coefficients in Table 4. Tree removal is in fraction of 1, i.e. a removal of 0.4 signifies 40% of trees removed.

Since both N leaching and N deposition decreased during the observation period a direct causal link between these two parameters seems obvious. This hypothesis was tested by extracting annual predictions for the variations in climate (potential evapotranspiration, minimum site water balance and leaching water), or N deposition from the regression model in Table 4. These estimates were compared with values predicted from the complete model and with observed data. This analysis revealed that changes of N deposition and climatic factors contributed equally to the changing N leaching rates (Figure 9).

**Figure 9:**
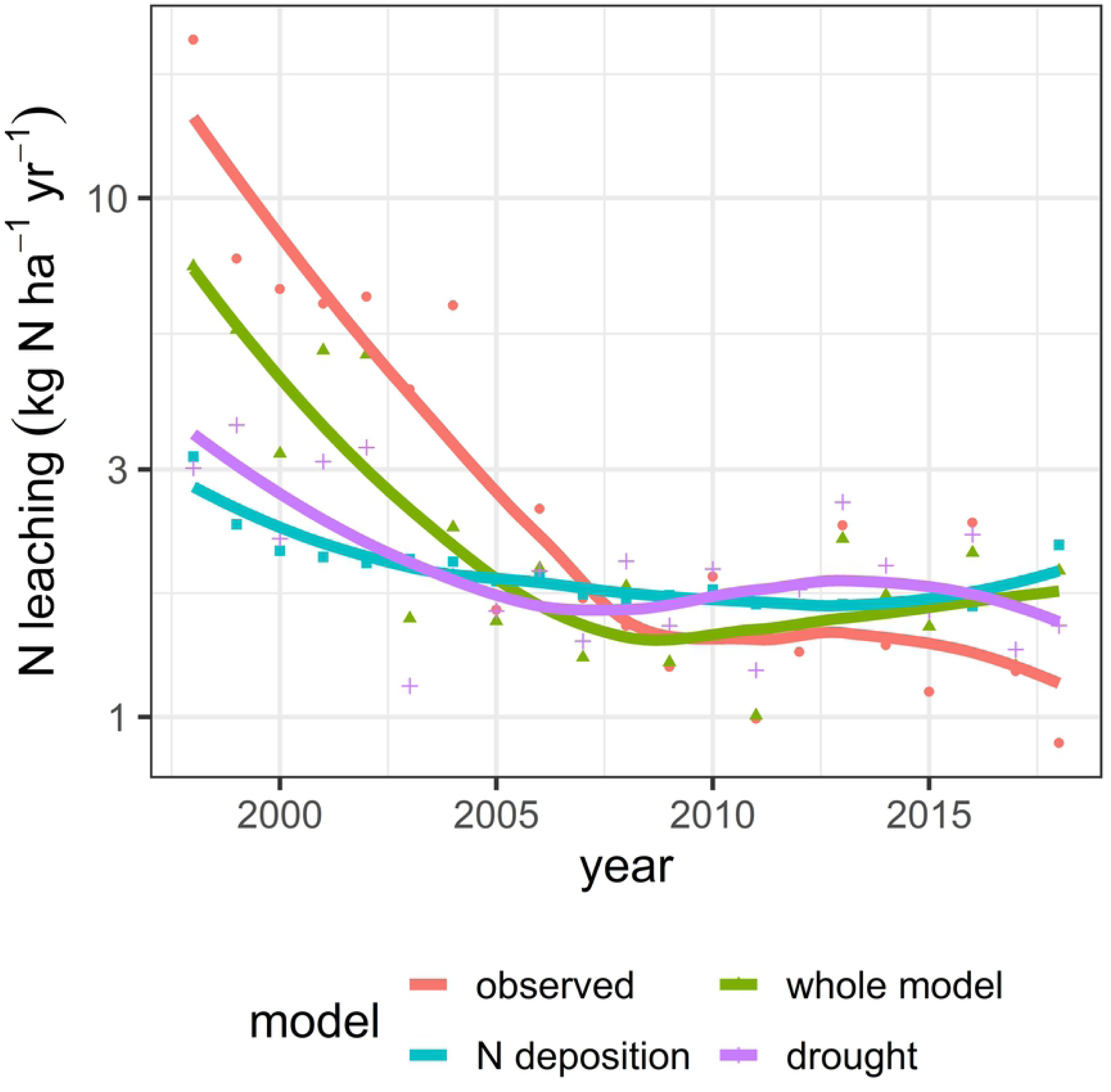
Predictions for the trend of the N leaching from the model in Table 4 for the complete model (“whole model”), for annual variations in precipitation and temperature (“drought”), and N deposition in comparison to the observed data (“observed”). The lines are a loess smoother, the points are annual means for predicted values or for observations.

Although N leaching in coniferous trees plots was clearly larger than in neighboring plots with deciduous trees (Figure_Supplementary 4), the effect of the proportion of coniferous trees on N leaching revealed not to be significant in the regression analysis. The problem with this analysis is that N deposition itself is also a function of tree species and is thus confounding.

## 4 Discussion

The results presented here suggest that soil acidification is still an issue in Swiss forests, both in terms of the extent and of the continuous progression in a large part of the plots. The thresholds observed for further in- or decrease of BC/Al ratio under the current deposition regime shown in Figure 5 can partially be explained by the soil chemical equilibria. The pH of 5.9 is quite close to the pH at which free CaCO3 disappears and the Ca buffer range passes over to the exchange buffer range (pH<6.2). Between pH 6.2 and 4.2 soils are buffered by the silicate and the exchange buffer ranges which have a lower capacity. At a pH 4.2 the aluminum buffer range is reached which has a high capacity (Ulrich 1988). No relation was found between N deposition and the speed of acidification in terms of decrease of BC/Al ratio. The results presented here confirm the effectiveness of the various buffer mechanisms,

N deposition decreased during the period of observation from 1998 to 2018, which seems to be the obvious reason for the observed decrease in N leaching. However, the statistical analysis revealed that climate, especially potential evapotranspiration and runoff, plays an equally important role for this trend. Analysis of the variation of N leaching in time showed also a significant contribution of the relative mortality or tree removal which can be detected in the three years following the event. Such an increase has been also observed after clearcut in a catchment ([20]) or after tree cut ([47]). The C:N ratio was not a significant predictor for N leaching. This is in contradiction to other studies ([18]) but can be explained by the low C:N ratio of the soils examined. 197 of 212 monitoring plots have a C:N ratio of <25 which was assigned by Gundersen as threshold for an enhanced leaching risk.

The higher N leaching under Norway spruce in comparison to beech observed on site pairs is in accordance to observations in Germany ([48], [49]). This may be explained by the higher N deposition in Norway spruce stands due to the higher leaf area, the higher surface roughness and the evergreen needles. In the overall data analysis the variable tree species was confounded with N deposition as tree species is an input to deposition modelling.

The present results do not allow conclusions on appropriate indicators of N eutrophication. Generally accepted indicators for eutrophication are increased N leaching, increased nitrate concentration in the soil solution, decreased C:N ratio of the forest floor or increased N in foliage ([50], [10]). In the monitoring plots presented in this study, foliar N concentrations in beech leaves are no longer correlated with N deposition while they were in the 1980’s ([21]), thus questioning foliar N concentration as a general indicator. C:N ratios in the forest floor are at a very low level at almost all sites so they allow no differentiation. While a large number of plots show increased N leaching at high N inputs, three plots have no N leaching and almost no detectable nitrate concentrations in the soil solution despite of a very high N input. None of the eutrophication indicators mentioned above apply for these plots.

## 5 Conclusions

The soil solution measurements have shown an increase in acidification in the majority of the sites between 2005 and 2018, even in base rich soils. The progression of acidification depends on the chemical state of a soil reflected in the buffer ranges: strongly acidified soils are in the aluminum buffer range and thus less susceptible to further changes. Driver of the observed acidification is the high N deposition resulting in high nitrate leaching and thus cation loss. In nutrient balance calculations, these cation leaching losses were the most important contribution to the balance, exceeding the input by weathering and deposition in a large number of plots and thus endangering forest sustainability ([15]).

The soil acidification has consequences for forest health. In the present network of forest monitoring plots, soil acidification effects have been reported in earlier studies. An increased risk of windthrow has been observed on soils with a base saturation of <40% ([51]). A decreased rooting depth was found on soils with a base saturation of <20% ([14]). Based on the relation between BC/Al ratio and base saturation presented in the current study, these base saturation thresholds translate to a BC/Al ratio in soil solution of 51 and 12, respectively. The BC/Al ratio of 12 is close to the ratio of 10 found by [52] to come to realistic estimates between BS, pH and BC/Al ratio in mineral soils.

High N deposition is still affecting most of the observed plots although air pollution measures have resulted in a decrease since the 1980’s. The current study provides information to predict the effect of climate change on leaching losses. It also quantifies the effect of forest management and tree mortality on the variation of N leaching in time.

## 6 Acknowledgements

The author acknowledges the financial support by the Federal Office for the Environment and by the cantons AG, BE, BL, BS, FR, SO, TG, ZH and the environmental offices of the cantons in Central Switzerland. I would also thank to the field and lab team of the Institute for Applied Plant Biology who performed the monthly sampling and analysis with a lot of patience and care: Delphine Antoni, Dieter Bader, Ute Schröder, Moïse Groelly, Caroline Stritt und Roland Woëffray. Part of the data analysis on N leaching was performed in a project on “Forest and Climate” supported by the Federal Office for the Environment in cooperation with Peter Waldner, WSL.

